# Cued reactivation during slow-wave sleep induces connectivity changes related to memory stabilization

**DOI:** 10.1101/185611

**Authors:** Ruud M.W.J. Berkers, Matthias Ekman, Eelco V. van Dongen, Atsuko Takashima, Marcus Barth, Ken A. Paller, Guillen Fernandez

## Abstract

Memory reprocessing following acquisition enhances memory consolidation. Specifically, neural activity during encoding is thought to be ‘replayed’ during subsequent slow-wave sleep (SWS). This natural tendency of memory replay can be induced by external cueing, known as “targeted memory reactivation”. Here, we analyzed data from a published study (van Dongen, Takashima, et al. 2012), where auditory cues reactivated learned visual object-location memories during SWS. Memory replay during sleep presumably involves a shift in connectivity across the brain. Therefore, we characterized the effects of memory reactivation on brain network connectivity using graph-theory. We found that cue presentation during SWS introduced increased network integration of the occipital cortex, a visual region that was also active during the object retrieval task. Importantly, enhanced network integration of the occipital cortex showed a behavioural benefit and predicted overnight memory stabilization. Furthermore, occipital cortex displayed enhanced connectivity with mnemonic regions, namely the hippocampus, parahippocampal gyrus, thalamus and medial prefrontal cortex during cue versus control sound presentation. Finally, network integration of early occipital cortex during cueing in SWS was related to increased activation of the bilateral parahippocampal gyrus, a region involved in coding for spatial associative information, at the post-sleep test. Together, these results support a neural mechanism where cue-induced replay during sleep promotes memory consolidation by increased integration of task-relevant perceptual regions with mnemonic regions.

## Introduction

The neural representations of recently acquired declarative memories are thought to be selectively reactivated during ensuing rest periods, particularly during slow-wave sleep (SWS) (Diekelmann and Born 2010; Diekelmann et al. 2011). For instance, hippocampal place cells in rats that co-activated during active behavior were found to be reactivated in concert during ensuing SWS (Wilson and McNaughton 1994). Later studies found that this coordinated reactivation, also dubbed replay, extended to the perceptual cortex involved in the initial processing of the stimulus, such as the visual cortex (Ji and Wilson 2007). Furthermore, evidence of neural replay has also been found in the medial prefrontal cortex (Euston et al. 2007; Peyrache et al. 2009; Genzel and Battaglia 2017). These results demonstrate the occurrence of neural memory replay in rodents, and have been shown to be behaviorally relevant for the retention of associative memories (Dupret et al. 2010).

Systems-level consolidation theory posits that neural memory traces are initially hippocampus-dependent, and subsequently integrate into neocortical networks (Squire et al. 1984; McClelland et al. 1995; Nadel and Moscovitch 1997; Frankland and Bontempi 2005). The neural trace is presumed to initially involve the hippocampus, which acts like a hub binding elements into a coherent memory trace, and the cortical regions that were involved in the initial processing of the stimulus at encoding (Nyberg et al. 2000; Polyn et al. 2005; Danker and Anderson 2010). During sleep, especially when the brain is in a highly synchronous state (i.e. SWS stage), specific tightly coupled oscillatory patterns are observed, including slow-oscillations, thalamic spindles, and hippocampal ripples (Siapas and Wilson 1998; Mölle et al. 2002, 2006; Staresina et al. 2015) that have been related to memory consolidation (Schabus et al. 2004; Marshall et al. 2006; Axmacher et al. 2008; Girardeau et al. 2009; Cox et al. 2012). Thus, it is plausible to posit that memory reactivation during sleep facilitates information exchange between the neocortex and hippocampus, enabling the strengthening of connections between these areas as well as amongst neocortical modules, thereby contributing to systems-level memory consolidation (Marshall and Born 2007).

Neural memory reactivation is difficult to observe in humans, as its timing is generally unknown. However, reactivation can also be induced using cues associated with information learned previously, termed “targeted memory reactivation”. For example, presenting an odor during SWS induces activation of the hippocampus and stabilizes associated memory traces (Rasch et al. 2007; Diekelmann et al. 2010, 2011). Furthermore, whereas studies have found that audio-visual-spatial associations can be stabilized in a targeted manner using specific auditory cues (Rudoy et al. 2009; Creery et al. 2015), these cueing effects can also affect all associations acquired within the same learning context (Oudiette et al. 2013). Van Dongen and colleagues (2012) performed targeted memory reactivation in the magnetic resonance (MR) scanner using auditory cues that were paired with specific object-location associations. Even though no consistent effect of cueing on memory stabilization was found, individual differences in the effect of cueing positively correlated with activity in the hippocampus, thalamus and cerebellum during SWS. Given repeated earlier findings of memory improvement from auditory cues presented during SWS (Oudiette et al. 2013; Ong et al. 2016; Schouten et al. 2017), it is likely that the noisy scanner environment contributed to excessive variability here, with some individuals effectively tuning out all auditory input, or alternatively that cueing affected all associations acquired within the same learning context (Oudiette et al. 2013).

Here, we re-analyzed data acquired by van Dongen and colleagues (2012) using a principled graph theoretical analysis approach to characterize whole-brain connectivity changes. Triggering reactivation of memory traces with auditory cues, could induce a process akin to spontaneous replay, which likely involves whole-brain connectivity changes facilitated by synchronous brain states during SWS. Conventional analysis methods are often unable to capture the coordinated whole-brain connectivity changes expected during memory replay. Thus, a network perspective can be leveraged using methods developed within the framework of graph theory (Bullmore and Sporns 2009). The *participation coefficient* is a metric that captures the extent to which a region integrates and distributes information as a result of the number and positioning of their contacts in the network (Power et al. 2013). Specifically, it quantifies network integration by looking at the importance of a given node for interactions between subnetworks. The participation coefficient can be used to index the integration of a given voxel (node) in the wider brain (network) during cued reactivation of visuospatial associations during SWS.

If a cue induces memory reactivation akin to replay, and this indeed plays a role in memory consolidation involving a hippocampal-neocortical trace shift, then one should expect a coordinated neural activation of multiple regions, including the sensory features of the cued memory (in this case visual object features in the occipital cortex (DeYoe et al. 1996) and the spatial layout in the parahippocampal gyrus (Aminoff et al. 2007)), as well as the key mnemonic regions (such as the hippocampus, thalamus, and medial prefrontal cortex). This increased integration of these and other regions would be expected to be beneficial for memory stabilization across sleep and result in an increased involvement of the neocortex in the memory trace.

## Materials and Methods

The Materials and Methods have been described extensively in the original report (van Dongen, Takashima, et al. 2012). Here, we summarize relevant details about the experimental setup, supplemented with details about the data analysis performed for this report.

### Participants

This re-analysis includes data from all 22 participants included in the original report (for complete description, see van Dongen et al., 2012). Briefly, 56 participants were initially recruited from the online research participant system of the Radboud University Nijmegen (age range 18–27; 14 males). Twenty-two participants in the sample reached a sufficient amount of SWS in the scanner to allow at least 80 % of the cueing protocol to be completed whilst in the SWS stage. Furthermore, these participants did not display micro-arousals in response to sound presentations, nor did they report explicit knowledge of the sounds having been presented while they slept. The experiment was conducted in accordance with national legislation for the protection of human volunteers in nonclinical research settings and the Helsinki Declaration and approved by the local ethics committee. Participants were compensated for participation with course credits or a monetary fee.

### Procedures

The experiment started between 7 and 8 PM, and participants went to sleep between 10 PM and midnight (see figure 1). Participants were placed in the MR scanner, and the sound volume was individually calibrated to a level at which the participant could distinguish individual sounds while the scanner was operating. Participants learned 50 object-location associations, each object-location pair being presented together with a unique sound associated with the object, inside the scanner. When all associations were learned to criterion (all objects placed within 4 cm from the correct location on two subsequent rounds), a pre-test was performed on all associations (Test 1). Next, participants were prepared outside the scanner for polysomnographic recordings, and placed in the scanner again for a 2 hr rest/sleep period, while electroencephalograph (EEG) was monitored online and fMRI data was continuously acquired. When participants had entered stable SWS for a period of time as visible on the ongoing polysomnography, sounds were presented to the participants using an MR-compatible headphone. Specifically, half of the sounds that were associated with specific object-location associations during learning were presented as ‘cue sounds’ during sleep (25 sounds presented twice, for a total of 50 cue sound presentations). Moreover, control sounds were presented that had not been previously associated with learning materials (5 sounds presented five times, for a total of 25 control sound presentations). After a 2 hr sleep, participants were awoken and subsequently were tested on all object-location associations (Test 2).

**Figure 1.**
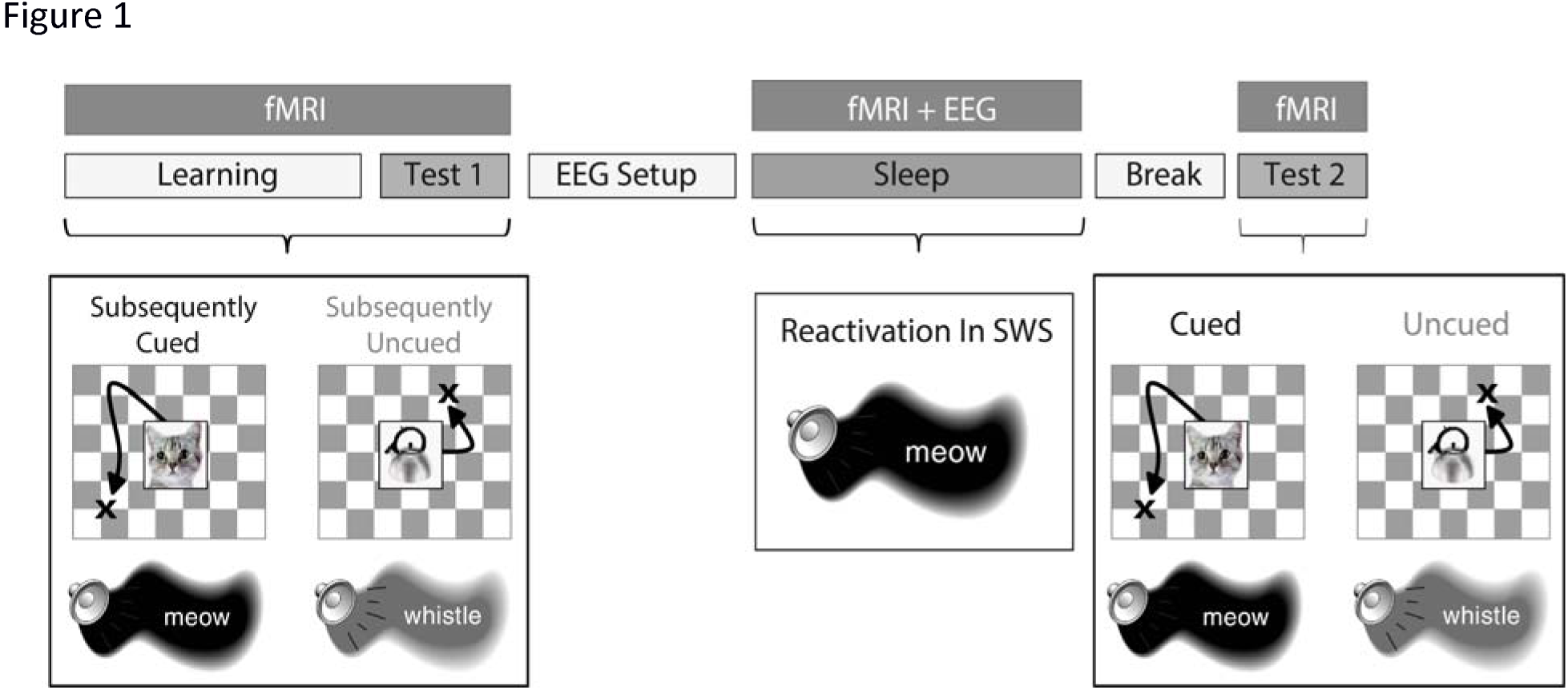
Schematic depiction of the experimental procedure. *Participants learned 50 object-location associations, presented simultaneously with object-related sounds, inside the MR-scanner. After a baseline test (Test 1), participants were set up for polysomnographic recordings and went to sleep inside the MR-scanner. Half of the learned associations were cued with sounds during slow-wave sleep (SWS). After awaking and a short break, participants performed another post-sleep test (Test 2). Figure adapted with permission from the initial report (van Dongen et al., 2012)*.

### Object-location test

The object-location task was performed in three stages: learning, Test 1 and Test 2. Here, we focus on the behavioral data from Test 1 and Test 2 for further analysis. Participants learned the location of 50 object pictures. For each participant, 50 objects were randomly assigned to 50 screen locations (screen size: 47 × 35 cm, resolution: 1024 × 768, viewing distance: 60 cm). In the learning phase, participants first passively viewed all 50 objects in their respective screen locations (duration: 3s presentation followed by a 1s inter-stimulus interval), paired with hearing the object related sound (e.g. a cat’s ‘meow’ when the object was a cat, sound duration: 500 ms). Next, the learning phase continued with several iterative rounds of active learning, where an object was presented at the center of the screen simultaneously with the auditory stimulus. Participants were then required to place the object in its original location using an MR-compatible joystick, and to press a button to confirm the object placement (self-paced timing). Feedback was given in every round of learning by displaying the object in the correct location for 3s. Objects were presented in a random order, and with every new round those objects were excluded that had been placed within 4 cm from the correct location on the two preceding rounds. Upon reaching criterion performance on all objects, participants performed the pre-sleep test (Test 1) for all objects without feedback. An identical post-sleep test was performed after sleep (Test 2). During tests, object placement was followed by an inter-stimulus period of fixation (duration jittered between 3 and 5s).

### MRI Data Acquisition

Participants were scanned using a reduced-noise Echo-Planar Imaging (EPI) sequence with sinusoidal gradients to avoid acoustic resonances of the scanner (Schmitter et al. 2008). Functional (T2*) images were acquired with whole-brain coverage (28 axial slices, ascending slice acquisition, repetition time (TR) = 2,511 ms, echo time (TE) = 38 ms, 90° flip angle, matrix = 64 x 64, bandwidth = 1,502 Hz per voxel, slice thickness = 3.5 mm, slice gap = 15%, field of view (FOV) = 244 mm). Structural (T1) images were acquired with a magnetization-prepared rapid acquisition gradient echo sequence (176 sagittal slices, TR = 2,250 ms, TE = 2.95 ms, 15° flip angle, matrix = 256 x 256, slice thickness = 1.0 mm, FOV = 256 mm).

### MRI Data Preprocessing

The fMRI data acquired during Test 1, the sleeping period and Test 2 were preprocessed using standard routines implemented in SPM8 (www.fil.ion.ucl.ac.uk). The first five volumes of each functional EPI run were discarded, and an outlier algorithm was used to check for corrupted slices from each image and replaced using between-volume interpolation. The functional images were then realigned, and coregistered to the structural image, spatially normalized to the Montreal Neurological Institute (MNI) EPI template (resampled at voxel size 2 × 2 × 2 mm), and smoothed using a Gaussian kernel (8 mm full-width at half maximum, FWHM). The structural images were segmented into gray matter, white matter, cerebrospinal fluid, and residual compartments (outside brain and skull) using the unified segmentation algorithm as implemented in SPM8. These compartments were used as masks to extract the mean intensity level across the whole time-series, and entered as compartment regressors to account for effects related to non-specific signal fluctuations.

### Network analysis during cueing

We employed a graph theoretic framework (Bullmore and Sporns 2009; Ekman et al. 2012) to analyze connectivity patterns during the presentation of sound cues in the scanned sleep period. The aim was to measure dynamic changes in the whole-brain network properties during memory reactivation, and specifically changes in integration of certain brain regions with the whole brain network. The metric of interest was defined as the participation coefficient (Guimera et al. 2005; Power et al. 2013; Backus et al. 2016), which quantifies for each node (i.e. voxel) the diversity of its inter-modular connections (Rubinov and Sporns 2010), specified as the amount of connections with nodes in other submodules relative to the total amount of connections. It therefore is a measure of the importance of a given node for global intermodular integration across the brain.

A high-pass filter with a cutoff period of 128s was applied to the functional time series. Sound presentation during sleep was modelled using a separate regressor for each sound presentation (to obtain trial-specific beta-estimates for the 50 cue sound presentations and 25 control sound presentations) with a duration of 5 seconds each. The design matrix also included six regressors of no interest to account for head movement, and the three compartment regressors to account for non-specific signal fluctuations. The resulting beta estimates were concatenated separately for cue sound and control sound trials. This procedure resulted in two separate beta time-series (i.e., cue sounds and control sounds) for all voxels in the brain. Next, voxel-wise correlation coefficients of beta time-series were computed to quantify pairwise functional connectivity (Rissman et al. 2004) for each condition separately (i.e., cue sounds and control sounds). The connectivity matrices were thresholded by preserving only significant connections (*p* < 0.05 False discovery rate (FDR) corrected) and setting all other connections, as well as all negative correlations to zero. Following the procedure used by Power and co-workers (2011), modular network structure was first derived across all sound presentations using a connectivity matrix based on the 116 anatomical regions defined by the AAL (Automated Anatomical Labeling) atlas (Tzourio-Mazoyer et al. 2002). Here, beta time-series were averaged across voxels of the region, and a 116 × 116 region-by-region connectivity matrix was constructed. After thresholding this connectivity matrix (edges > 0, p < 0.05 FDR corrected), the matrix was parcellated into subnetworks using modularity detection according to the Louvain method (Blondel et al. 2008). Next, each voxel in the whole-brain voxel-wise connectivity matrix was assigned to a module, allowing the calculation of their participation coefficient ranging from 0 (provincial hub: connections are only present within its module) and 1 (connector hub: connections are only present with other modules; Power et al., 2013). The resulting participation coefficient maps for cue sounds and control sounds were contrasted on an individual subject level and this contrast was subsequently tested at the second level using a one-sample t-test.

### Psychophysiological interaction (PPI) analysis during cueing

The previous analysis determined which of the regions increased in global inter-modular network integration. To determine which regions in other modules of the brain were connected with the ‘hub’ (the region found to display increased network integration during cueing), we performed a follow-up seed-based functional connectivity analysis. This psychophysiological interaction (PPI) analysis served to determine which brain regions were more connected with the hub during the presentation of cue versus control sounds. Therefore, we extracted the physiological signal time series from the seed region found in the network analysis (cluster-forming threshold at *Z* > 2.33) to serve as a physiological regressor. This was entered into a general linear model (GLM) along with the psychological variable (cue versus control sounds) and the psychophysiological interaction regressor. Small-volume correction was used for the bilateral parahippocampal gyri, as this region was activated by the object-location retrieval task (van Dongen, Takashima, et al. 2012), and the thalamus and hippocampus, as these are smaller structures hypothesized to be involved in memory reactivation. Here, anatomical masks implemented in the IBASPM 71 Atlas of the WFU Pick Atlas Tool in SPM were used (Iturria-Medina et al. 2007).

### Activity analysis during pre-and post-sleep test

We next assessed whether network integration of visual cortex during cueing in SWS results in increased involvement of cortical associative regions at retrieval. For this reason, we contrasted the neural activations observed during the pre-sleep (Test 1) and post-sleep (Test 2) tests, and related this contrast to the participation coefficient of the occipital cortex as observed in the graph theory analysis (see Results section). The fMRI data for Test 1 and Test 2 were analyzed in one GLM. Similar to van Dongen and colleagues (2012), this model included regressors for cued and uncued associations, which were modeled as delta functions to be convolved with the canonical hemodynamic response function (HRF), along with temporal derivatives provided by SPM8. The design matrix included six regressors of no interest per test session to account for head movement using the realignment parameters, as well as compartment regressors accounting for non-specific signal fluctuations. Furthermore, a high-pass filter with a cutoff period of 128s was implemented to remove low-frequency fluctuations from the time series. Parameter images were generated on the first level (for each individual participant) based on a contrast between all trials of Test 1 and all trials of Test 2, to assess changes in retrieval activity from the pre-to the post-sleep test. We collapsed across cued and uncued associations because network participation of the occipital cortex was not specifically related to only the cued associations in the learned set (see ‘Results’). These contrast images were entered into a one-sample t-test on the second level, with the participation coefficient of the occipital cortex during cue versus control sound presentation entered as a covariate. Small-volume correction was used for the bilateral parahippocampal gyri, as this region was activated by the object-location retrieval task and cue-sound presentation (van Dongen, Takashima, et al. 2012). Here, an anatomical mask of the parahippocampal gyrus implemented in the IBASPM 71 Atlas of the WFU Pick Atlas Tool in SPM was used (Iturria-Medina et al. 2007).

## Results

### Network changes following cueing

We probed changes in network dynamics in response to the presentation of cue sounds versus control sounds. The presentation of cue sounds was hypothesized to induce reactivation of the corresponding memory trace, which would presumably be reflected in increased integration of areas that represent sensory features of the cued memory representation, e.g. occipital cortex. Indeed, we found a specific increase in the participation coefficient of early visual (occipital) cortex (peak MNI coordinates: *x,y,z* = [4, −70, 6], *z*_*21*_ = 3.12, cluster-forming threshold at Z>2.33, corrected for the whole brain at p < 0.05 using Gaussian Random Field Theory), indicating that occipital cortex displays an increased inter-modular connectivity when presented with cue sounds during deep sleep. The region observed overlaps with regions that were activated in response to the object-location task during Test 1 (see figure 2b). This overlap may be indicative of replay of the original visuo-spatial neural memory trace in response to auditory cues presented during sleep, in the absence of any external visual input.

**Figure 2.**
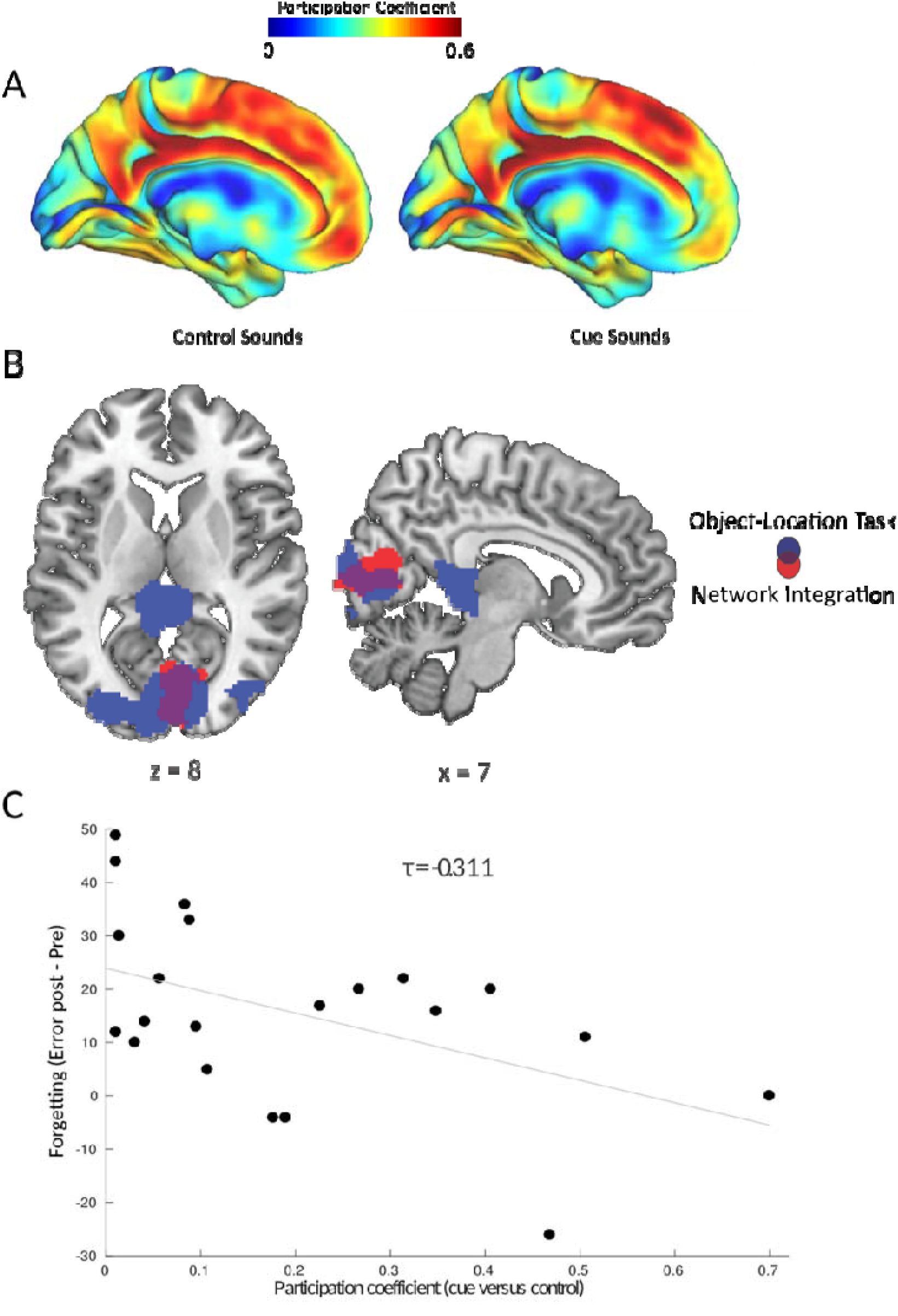
Replay-related changes in network integration predict behavioral memory stabilization. *A) Participation Coefficient mapped for every voxel in the brain during the presentation of control sounds (not previously paired with object-location information, left panel) and cue sounds (previously paired with object-location associations, right panel)*. *B) Increased network integration was found only in the occipital cortex during the presentation of cue sounds versus control sounds during SWS. This region overlapped with the set of regions activated in response to the object-location association task. Parametric maps were superimposed onto a template brain, using the cluster-forming threshold of *Z*>2.33*. *C) The increase in participation coefficient of the occipital cortex during cueing predicted memory stabilization, indicated by reduced forgetting (expressed as the difference in error distances) between test 1 (pre) and test 2 (post)*.

To determine what brain regions this area in the occipital cortex preferentially connects to when recruited in the network by the auditory cues, we performed a psychophysiological connectivity analysis. The seed was defined as the region in cortex that displayed an increase in participation coefficient during cueing (cluster-forming threshold at *z* > 2.33). The task contrast was formed by the presentation of cue sounds versus control sounds. The occipital cortex was found to be connected to the left hippocampus (peak MNI coordinates: *x,y,z* = [-32, −16, −18], *z*_*21*_ = 3.36), right hippocampus (peak MNI coordinates: *x,y,z* = [22, −2, −22], *z*_*21*_ = 2.98), left parahippocampal gyrus (peak MNI coordinates: *x,y,z* = [-26, −22, −26], *z*_*21*_ = 3.10), left thalamus (peak MNI coordinates: *x,y,z* = [-4, −2, −0], *z*_*21*_ = 3.11), and right thalamus (peak MNI coordinates: *x,y,z* = [2, −16, −12], *z*_*21*_ = 2.67), and a medial prefrontal region extending from premotor cortex to anterior cingulate cortex and into dorsomedial prefrontal cortex (see figure 3, all cluster-forming threshold at *z* > 2.33, cluster-size corrected for the whole brain, or reduced search regions based on predefined anatomical areas for the bilateral hippocampus, parahippocampal gyrus and thalamus, thresholded at p < 0.05 using Gaussian Random Field Theory). Thus, during cueing, the occipital cortex was connected to several regions where memory replay has been shown to occur (the hippocampus, medial prefrontal cortex), and that are involved in oscillatory patterns of activity (spindles, ripples, slow-waves) that allow for systematic interactions between brain regions (e.g., the hippocampus, medial prefrontal cortex and thalamus).

**Figure 3.**
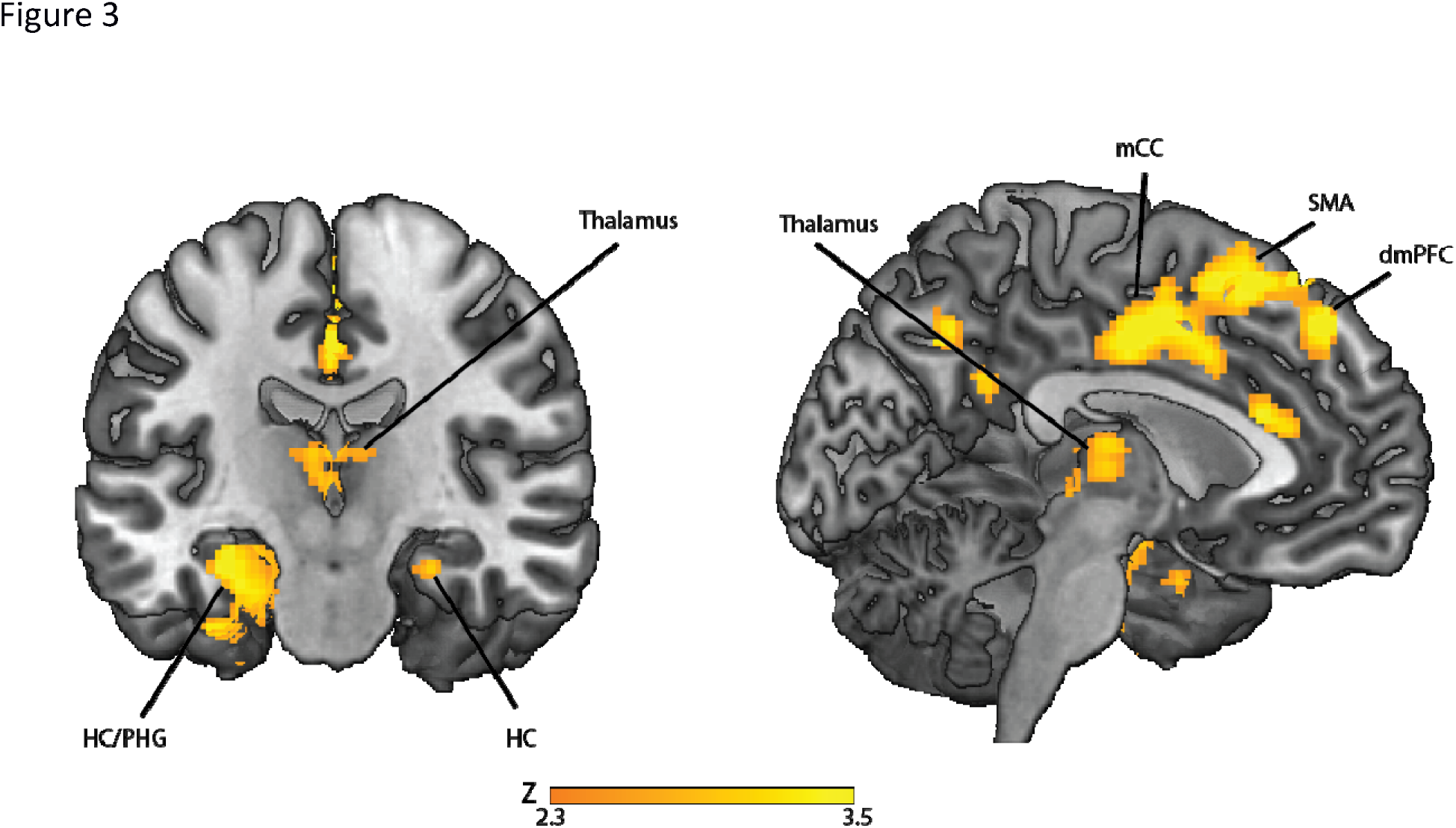
Regions displaying greater coupling with occipital cortex during cueing in SWS. *Regions displaying stronger coupling with the occipital cortex for cue sounds compared to control sounds. Parametric maps were superimposed onto a template brain, using the cluster-forming threshold of Z>2.33. HC = Hippocampus, PHG = Parahippocampal Gyrus, mCC = Middle Cingulate Cortex, SMA = Supplementary Motor Area, dmPFC = Dorsomedial Prefrontal Cortex*.

### Network dynamics related to behavioral cueing effect

As reported earlier (van Dongen, et al. 2012), memory performance decreased on average (Test 1 performance: error = 2.74 ± 0.12 cm; Test 2 performance: error = 3.12 ± 0.14: Δerror = 0.37 ± 0.08; t_21_ = 4.48, p < 0.001), both for cued (Δerror = −0.44 ± 0.11 cm, *t*_*21*_(3.86, *P* = 0.001)), and uncued associations (Δerror = −0.33 ± 0.11 cm, *t*_*21*_(2.56, *P* = 0.018)). There was no effect of cueing on participants’ memory accuracy as tested after sleep (*F*_1,21_ = 0.83; *P* = 0.374). Thus, there was forgetting across the sleep/rest session, and this forgetting appeared to be unspecific across cued and uncued object-location associations.

However, there was substantial variability in the effect of cueing on retention. Therefore, different neural responses to cueing might explain individual differences in the effect of cueing. Particularly, increased network participation of the occipital cortex could be an index of how much the visual-spatial memory trace became reactivated in a given subject when auditory cues were presented. Indeed, the increased participation coefficient during cue sound presentation was found to be associated (see figure 2) with reduced overnight forgetting of visual-spatial object locations (τ_21_ = −0.311, p = 0.045). This relationship was specifically present for the visual-spatial locations that were associated with sound cues (τ_21_ = −0.338, p = 0.029), but not for other locations (τ_21_ = −0.211, p = 0.174), although the difference in these correlations did not reach significance (Williams-Hotelling test, t = 0.433, p = 0.670).

The distribution of the increase in participation coefficient followed a non-normal positively skewed distribution, with about half of participants displaying only a minimal increase in participation coefficient during cueing (PC < 0.1). Therefore, we performed a median split to better characterize the group that experienced an increase in network participation of the occipital cortex (PC+), versus the group with no increase (PC=). The PC= group displayed significant forgetting (Δerror = 0.68 ± 0.32; t_10_ = 6.042, p < 0.001), for both cued (Δerror = 0.69 ± 0.41; t_10_ = 5.620, p < 0.001) and uncued (Δerror = 0.48 ± 0.44; t_10_ = 3.636, p = 0.005) associations. In contrast, the PC+ group did not display any forgetting (Δerror = 0.16 ± 0.35; t_10_ = 1.551, p = 0.152), for both cued (Δerror = 0.20 ± 0.57; t_10_ = 1.186, p = 0.263) and uncued (Δerror = 0.12 ± 0.61; t_10_ = 0.653, p = 0.529) associations. Note that we do not report the statistical test to compare performance differences between the two groups, as this comparison would not be statistically independent from the across-subject correlation reported earlier. This result indicates that in the group where there was an increased network participation of occipital cortex, overnight forgetting was reduced, but the effect was not specific to the cued items, extending to other object-location associations encoded within the same spatial layout and temporal learning context (see figure 4).

**Figure 4.**
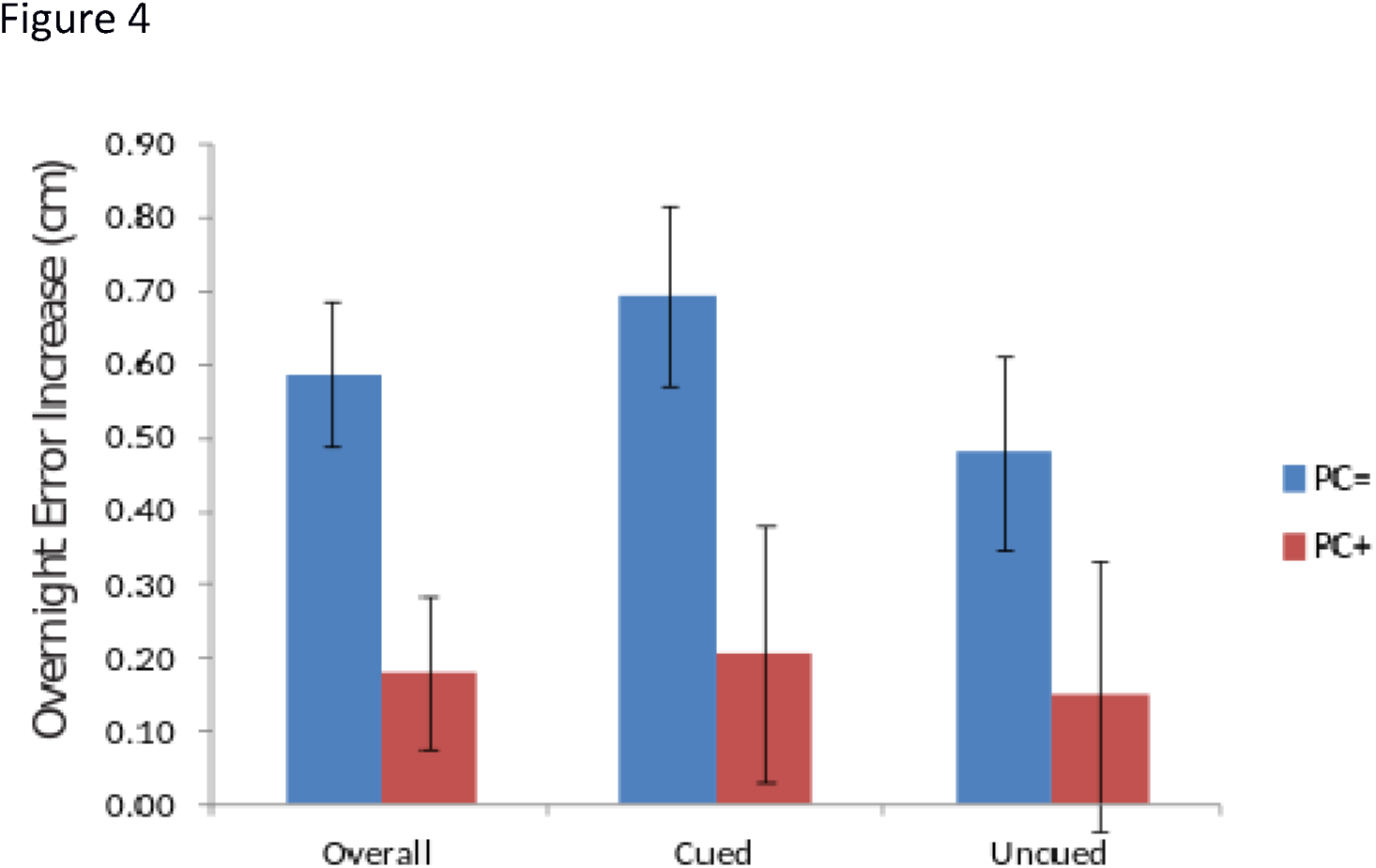
Overnight forgetting, split by network integration in occipital cortex. *The bar graph depicts the change in error distance between the pre-sleep Test 1 and the post-sleep Test 2 for a group where there was an increase in network integration (as measured with the participation coefficient) in occipital cortex (PC+) and a group where no such increase was found (PC=). The groups were defined using a median split on the participation coefficient values acquired from the occipital cortex in the cue sounds > control sounds comparison. The error distance is as the distance between the location where the object is placed at test, and the actual object location as presented during the learning phase. A higher overnight error increase indicates a reduction in accuracy of placement across sleep, indicative of increased forgetting. The group where cueing induced increasing network integration in the occipital cortex (PC+) displayed reduced forgetting overall, indicating that, in this group, cueing stabilizes memories overnight*.

### Overnight activity changes related to network integration during cueing

We then considered whether the increased network integration of the occipital cortex during sleep had subsequent effects on neural activity when retrieving object-locations during the post-sleep test. Here, we found that the participation coefficient was related to a positive increase in retrieval-related activity from Test 1 to Test 2 in both the posterior left parahippocampal gyrus (peak MNI coordinates: *x,y,z* = [-16, −32, −18], *z*_21_ = 4.14), and posterior right parahippocampal gyrus (peak MNI coordinates: *x,y,z* = [16, −32, −18], *z*_*21*_ = 3.26, cluster-forming threshold at Z>2.33, corrected for a reduced search volume based on anatomical masks of the left and right parahippocampal gyrus at a threshold of p < 0.05 using Gaussian Random Field Theory). Thus, increased network integration of the occipital cortex during cueing had a subsequent effect on the neural regions activated during the retrieval of object-locations at Test 2, thereby increasingly recruiting the bilateral parahippocampal gyrus.

## Discussion

Here, we investigated the whole-brain neural network reorganization induced by cued reactivation of memory traces during deep sleep, and its behavioral and neural effects on the subsequent retrieval of those memories. For that purpose, we re-analyzed fMRI data reported in the study by van Dongen and colleagues (2012) using whole-brain connectivity measures. We demonstrate that when participants were in a state of slow-wave sleep (SWS), absent of visual input, and were cued with sounds associated with previously learned visuospatial information, the occipital cortex displayed increased network integration as measured with the participation coefficient. Furthermore, subjects who displayed an increase in cue-induced network integration of the occipital cortex showed increased memory stabilization. This finding is indicative of a process in which the sound cue induces a global reprocessing of the learning episode, akin to memory replay, which is conducive to subsequent memory retention. Note that while the occipital cortex is not directly related to mnemonic processes, previous studies have shown that cue-triggered reactivation can lead to instantiations of previously experienced episodes in this region (Xu et al. 2012; Ekman et al. 2017). It is likely that this reinstitution is coordinated with other mnemonic regions. Indeed, various regions displayed increased connectivity with the occipital cortex during the presentation of cue sounds: memory and replay-related regions, such as the hippocampus, thalamus and medial prefrontal cortex; and higher-order associative cortices implicated in the encoding of the memory trace, such as the parahippocampal gyrus. These results are in line with a study by Ji & Wilson (2007) that shows coordinated replay between the hippocampus and a perceptual region that was involved in the initial processing of the stimulus (here: the occipital cortex). Cueing during SWS also showed an enduring neural effect at memory retrieval as measured during the post-sleep retrieval test. Specifically, subjects with greater network integration of the occipital cortex during SWS recruited the parahippocampal gyrus more at subsequent retrieval.

It should be noted that the network reorganization found during the presentation of cue sounds was not specifically related to memory stabilization of those object-location associations associated with the cues. The association of network participation with memory stabilization was numerically larger for cued, versus uncued material, but the interaction was not statistically significant. There could be various explanations for this unspecific effect. First, the MRI scanner is a suboptimal environment for auditory presentation during sleep. Therefore, it is conceivable that the acoustic stimuli presented during deep sleep in a noisy environment effectively modulated ongoing brain processes in some individuals, but not others, contributing to large inter-individual differences and lower power of demonstrating this effect than in other studies conducted outside of the MR-scanner environment (Rudoy et al. 2009). Indeed, we observed a non-normal positively skewed distribution of increase in participation coefficient during cueing. There was an overall effect on memory retention for participants who showed increased participation coefficient during cueing – but there did not appear to be a cue-specific memory benefit even when only considering this sub group. It is also possible that the study was underpowered to observe an interaction effect, and that a larger sample would have rendered a significant interaction. Furthermore, it is possible that the cueing had a non-specific effect on all material acquired within the same temporal and spatial learning context (as also suggested by the results reported in Oudiette et al., 2013). Indeed, all object-location associations were acquired within the same learning context; the same scanning session and temporal learning sequence, and most importantly, all objects were located on the same two-dimensional spatial grid. Therefore, it is plausible that the various object-locations became integrated into a unitary map, or spatial schema (Tse et al. 2007; van Buuren et al. 2014) at initial learning and/or during replay (Durrant et al. 2015; Hennies et al. 2016). As such, object locations could have been stored not only with reference to the two-dimensional grid, but also directly or indirectly in spatial reference to the other objects. As such, a sound cue could have reactivated certain object-location associations, but also objects located within the objects’ vicinity. In fact, it has been proposed that sleep-dependent memory replay is instrumental in the integration of memories into a cognitive schema (Lewis and Durrant 2011). Thus, while network integration of the occipital cortex during cueing is demonstrated to be positively related to overall memory stabilization of a set of object-location associations, the fact that an integrated spatial schema was cued may have contributed to non-specific reactivation of all object-locations within this set.

Bearing in mind this caveat, our results show a network reorganization in response to cues presented during SWS that is consistent with an active model of sleep-dependent memory consolidation. Previous research has shown that whole-brain connectivity patterns display a general reduction in thalamo-cortical and cortico-cortical connectivity and an increase in local clustering during SWS (Spoormaker et al. 2010). Against this backdrop, a hippocampal-neocortical dialogue takes place facilitating systems-level consolidation by integrating hippocampal-dependent memories into neocortical storage sites (Buzsáki 1996; Diekelmann and Born 2010; Mitra et al. 2016). This hippocampal-neocortical dialogue is orchestrated through oscillatory electrophysiological patterns characteristic of SWS and related to memory replay (Diba and Buzsáki 2007; Davidson et al. 2009). Specifically, slow (delta wave) oscillations propagate across the brain and to the medial temporal lobes, including the hippocampus, exerting a global control over spiking activity (Massimini et al., 2004; Nir et al., 2011). Mechanistically, the up-state of slow oscillations has been found to enable thalamic sleep spindles in the sigma band, which in turn cluster high-frequency hippocampal ripples in their troughs (Staresina et al., 2015). These hippocampal ripples then are proposed to allow for a precisely timed exchange of mnemonic information between the hippocampus and neocortex, potentiating and strengthening the neocortical memory trace. Indeed, the importance of slow oscillations and spindles for memory consolidation has been demonstrated with behavioral correlations (Schabus et al., 2004; Holz et al., 2012). Moreover, intracranial recordings in humans have found time-locked ripple events during sleep in both the hippocampus and neocortex, the latter being related to memory consolidation (Axmacher et al., 2008). Experimental disruption of hippocampal ripples in rats impaired consolidation of hippocampal-dependent spatial memories (Girardeau et al., 2009). Therefore, during SWS, hierarchically nested oscillations presumably serve to consolidate memories by tightly coupling the neocortex (especially medial prefrontal cortex), thalamus and medial temporal lobes, which are the regions that are displaying an increased interconnectivity during cueing here.

Thus, during SWS, there is increased cross-talk between the hippocampus, thalamus and neocortex. If external auditory cueing would induce a reactivation of the visuo-spatial memory traces, one would expect that cortical representation areas corresponding to the modality of the memory trace would be recruited to participate in this cross-talk. Indeed, the occipital cortex displayed an increase in global network integration during cueing, and when probing the neural connectivity changes paired with this increase, we found that this region increases its connectivity with the regions involved in active cross-talk during SWS, namely the hippocampus, thalamus and medial prefrontal cortex. These regions resemble those that are involved in the oscillatory patterns that orchestrate hippocampal-neocortical cross-talk (Mölle et al. 2006; Staresina et al. 2015). At first sight, it might be surprising that the auditory cortex is not recruited in response to auditory cueing. However, it should be noted that a contrast was made between two conditions in which sounds were presented. The critical difference in the contrast is that cue sounds were associated with specific visuo-spatial memories, but not control sounds. It remains the case that one could still expect increased connectivity of the auditory cortex with visual, spatial, and memory regions during cueing, the sounds being associated with the object-locations. As psychophysiological interaction analyses were informed by results of the graph theory analyses, we did not further probe connectivity from an anatomically defined seed in the auditory cortex here. Despite this caveat, it remains to be noted that cueing recruited the occipital cortex to engage with mnemonic regions that have been shown to display increased cross-talk during SWS, and this promoted the stabilization of the associated memory traces in neocortical storage sites.

Sleep supports memory retention in a selective manner, and information is prioritized based on perceived future relevance at encoding (Wilhelm et al. 2011; van Dongen, Thielen, et al. 2012) emotional salience (Hu et al. 2006; Wagner et al. 2006; Payne et al. 2007) and consistency across episodes and with prior knowledge (Tamminen et al. 2013; Durrant et al. 2015). Possibly, relevant information is selectively prioritized for active processing during sleep, whereas non-prioritized memory traces are forgotten, perhaps aided by homeostatic regulation of synaptic plasticity through global downscaling (Tononi and Cirelli 2003, 2006; Genzel et al. 2014). The net result is an increase in signal-to-noise for the prioritized memory traces in the neocortex, contributing to memory stabilization. Indeed, this appears to be the case in the parahippocampal gyrus, where an increase in retrieval-related activation of the posterior parahippocampal gyrus at post-sleep test versus pre-sleep test was related to an increase in network participation of the occipital cortex during cueing. The parahippocampal gyrus is known to be involved in representing the local visual environment (Epstein and Kanwisher 1998) and learning spatial layouts (Aguirre et al. 1996; Epstein et al. 1999). The increased involvement of the parahippocampal gyrus during post-sleep retrieval is consistent with a role of the parahippocampal gyrus in binding together low-level visual information from downstream occipital cortex. Notably, the parahippocampal gyrus was activated already during retrieval at the pre-sleep test, showed selective increased activation in response to cueing, and displayed increased connectivity with the occipital cortex during cued reactivation, as originally reported in van Dongen & colleagues (2012). The results reported here further elucidate the mechanisms involved in targeted memory reactivation by taking a network perspective.

In sum, we show here that inducing the reactivation of object-location memories with associated auditory cues during SWS increases the integration of occipital cortex with the global brain network, specifically those regions involved in memory replay, such as the medial temporal lobe, thalamus and medial prefrontal cortex. Furthermore, global network integration of the occipital cortex during cueing is predictive of overnight memory stabilization of learned materials and an increased involvement of the parahippocampal gyrus during the post-sleep retrieval. These findings highlight how graph theory analysis can be used to assess whole-brain connectivity patterns during targeted memory reactivation in SWS, and contribute to a better understanding of sleep-related memory consolidation.

**Figure 5.**
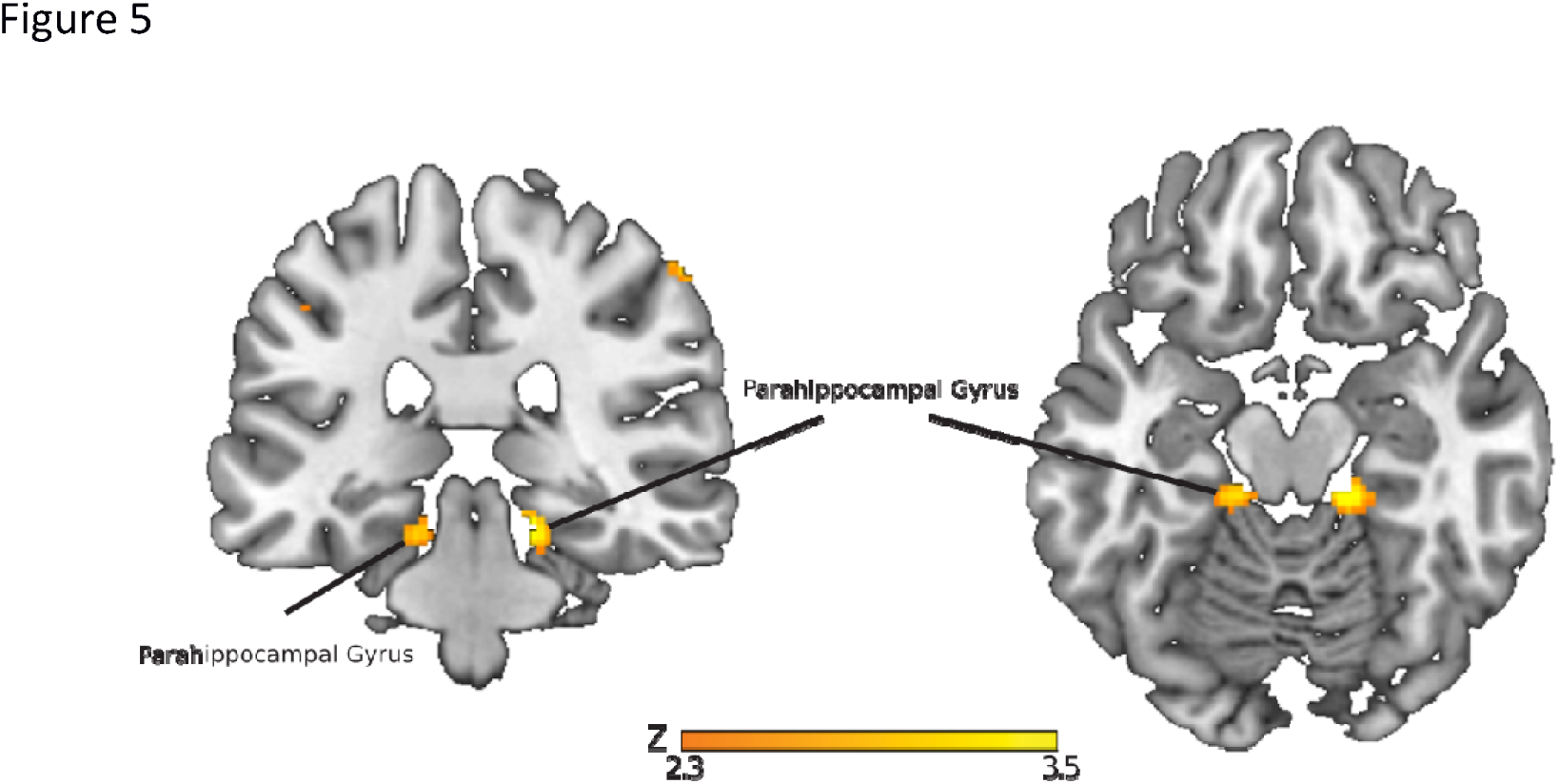
Overnight activity increases related to network integration of occipital cortex. *Regions where there is a relation between on the one hand, an activity increase between the pre-sleep and post-sleep test and, on the other hand, increased network integration of occipital cortex during cueing in the SWS between Test1 and Test 2. Parametric maps were superimposed onto a template brain, using the cluster-forming threshold of *Z*>2.33*.

## Acknowledgement

R.M.W.J.B., E.V.vD. and G.F were supported by a grant from the European Research Council (“Neuroschema” ERC R0001075). G.F. was also supported by a grant from the NWO (“Language in Interaction”). E.V.v.D (400-06-110) and A.T. (451-06-006) were supported by grants of the Dutch Organization for Scientific Research (NWO).

